# Waist rotation angle as an indicator of probable human collision avoidance direction for autonomous mobile robots

**DOI:** 10.1101/2022.05.03.490539

**Authors:** Hideki Tamura, Tatsuto Yamauchi, Shigeki Nakauchi, Tetsuto Minami

## Abstract

Currently, the probability of pedestrians on the streets of smart cities encountering autonomous mobile robots (AMRs) is increasing. Previous studies have discussed collision avoidance between humans as they cross paths. Their avoidance behavior toward AMRs, however, remains unclear. To address this, we experimentally investigated the avoidance direction a human would choose against an AMR approaching head-on. This experiment included recording human locomotion under such a scenario. Further, an AMR was programmed to approach from various starting points, including directly through the participants (head-on collision). The participants were directed to evade it by moving rightward or leftward, and their paths were tracked. We found that the participants did not strongly prefer either direction, suggesting that the avoidance direction is not solely determined by the participants’ attributes, such as their adherence to the traffic rules of their region. The probability of rightward evasion when the AMR approached head-on indicated that humans use different avoidance strategies when encountering other humans and obstacles. Moreover, the participants’ motion analysis revealed that they involuntarily twisted their waists in the avoidance direction before they evaded the AMR. These results suggest that this twist is the most important predictor of the avoidance direction. These findings could be encoded into the programs of AMRs to adapt these vehicles to our locomotory responses more organically.

## Introduction

In certain parts of world, the frequency of encounters with autonomous mobile robots (AMRs) on urban streets has been increasing, because the governments and industries in these regions are actively promoting developments toward smart cities (Golubchikov & Thornbush, 2020; Liu et al., 2020; O’Grady & O’Hare, 2012). AMRs move along planned delivery routes to conserve time. However, they must ensure the safety of pedestrians, even under complex situations (Meerhoff et al., 2018). Thus, several studies in the field of robotics engineering have utilized human-like behaviors to develop robots that are safer and friendlier to use (Kamezaki, 2019; Kruse et al., 2013; Silva et al., 2019; Tamura et al., 2010; Tomizawa & Shibata, 2016). For instance, Kamezaki verified control methods whereby a robot guided or made way for a human depending on the situation, based on the estimation of the positional relationship between the human and the robot (Kamezaki, 2019). This study reported that humans preferred to avoid a robot with a mutual concession, rather than either one entirely making way. These findings suggest the importance of such human-like control in robots, which positively influences humans’ impressions.

This participant must be approached from the perspective of cognitive science by testing the obstacle-avoidance techniques of humans. This knowledge could be applied to understanding human collision-avoidance behaviors when encountering robots. For example, humans employ suitable approaches to circumvent static obstacles such as barriers (Baxter & Warren, 2020; Fajen & Warren, 2003), a pole (Vallis & McFadyen, 2003) or two poles (Hackney et al., 2015, 2020). Other people, i.e., a type of obstacle, frequently walk into the participants’ surroundings, resulting in a dynamic environment (Fajen, 2013). In addition, a previous study discussed the techniques used by humans to avoid such dynamic obstacles approaching from an orthogonal direction (Basili et al., 2013; Olivier et al., 2013) and crossing paths (Bühler & Lamontagne, 2019; Souza Silva et al., 2018). In addition, several studies have replicated these real-world paradigms in virtual reality (Bühler & Lamontagne, 2019; Hackney et al., 2020; Lynch et al., 2018, 2021; Souza Silva et al., 2018) because the complete evasion of a possible physical collision and an increase in the number of variables of the experiment (such as adding human walkers in the environment) are beneficial for the participants.

However, extant literature includes few studies from the perspective of cognitive science on the evasive actions of humans against AMRs, which may become common dynamic obstacles in the future. Recent studies have focused on understanding human behaviors while walking with an AMR. For example, Vassallo et al. reported that humans made way for AMRs, even when they had the chance to pass first (Vassallo et al., 2017). In addition, their avoidance strategies varied depending on the target (human or AMR). Following this, the same group in another study programmed the AMR to replicate the human collision-avoidance strategy. The result revealed that humans responded to an oncoming AMR the same way they would to other approaching humans (Vassallo et al., 2018). These findings offer insights into the benefits of studies involving human locomotion and AMRs. However, these previous studies only examined situations wherein humans cross the AMR orthogonally. Thus, the technique used by humans to evade an AMR when forced to choose an avoidance direction for safety, such as during imminent head-on collision, remains unclear.

Therefore, this study aims to understand the avoidance strategy used by humans against AMRs approaching head-on. Specifically, we examine humans’ preference for a specific side (right or left). In this regard, Souza Silva et al. examined the method used by humans to evade other humans and a cylindrical object approaching head-on in a virtual environment (Souza Silva et al., 2018). Their findings revealed that humans predominantly avoided other humans by moving rightward (right: 75 % vs. left: 25 %); however, they did not report this bias in the case of the cylindrical object (55 % vs. 45 %). This suggests that humans change their evasive action depending on the target. Accordingly, we verified whether humans’ avoidance strategies for an AMR resembled those for other humans or objects.

We also examined the determination of the avoidance direction under such situations. A previous study suggested that humans exhibit a tendency to move rightward to avoid collisions, owing to a cognitive bias derived from the traffic rules associated with right-hand driving (Souza Silva et al., 2018). If this were true, however, people who live in countries that follow the left-hand driving tradition would behave differently. In this work, we assumed that humans would find it challenging to instinctively decide the avoidance direction in unusual situations, such as when an uncommon object is approaching them. Thus, we conceived the locomotory response of humans to an inbound AMR by computing their avoidance probability with the AMR as the obstacle. We hypothesized that humans predominantly avoided collisions by moving to a specific side when they consider the AMR as 1) a human and 2) an inorganic object.

We used an actual AMR as the obstacle and recruited human participants living in an area following left-hand driving. Specifically, we conducted an experiment to verify the technique used by humans to evade the AMR approaching from five directions, including head-on. Simultaneously, a motion-tracking system was used to estimate their walking trajectories. Furthermore, we verified the indices that could be derived from human locomotion to predict the avoidance direction.

## Methods

### Participants

Sixteen students (one female and 15 male participants; average age of 22.9 ± 0.7) from Toyohashi University of Technology participated in this experiment. All participants declared right-handedness; the average hand preference was 9.1±1.9, as assessed by the Flinders Handedness Survey (FLANDERS) questionnaire (Nicholls et al., 2013; Okubo et al., 2014). All experimental protocols were approved by the institutional review board of Toyohashi University of Technology for involving human participants in the experiments, following the Declaration of Helsinki. Written informed consent was obtained from all participants for the publication of their details.

### Apparatus

#### Control system for AMR

We used a custom-made wheeled platform (Mega Rover Ver 2.1, Vstone) as the AMR (**Figure 1a**). The dimensions of the AMR were 460 mm × 320 mm × 800 mm (length × width × height). The onboard computer of the AMR controlled two motors connected to the axles of its wheels. Another computer installed in the experiment chamber received signals from a tracking system (mentioned below) and transferred them to the onboard computer via Wi-Fi. The robot operating system (ROS) Melodic with Ubuntu 18.04 LTS was used to control both computers. The participants were unfamiliar with this AMR, which ensured bias-free results.

**Figure 1.**
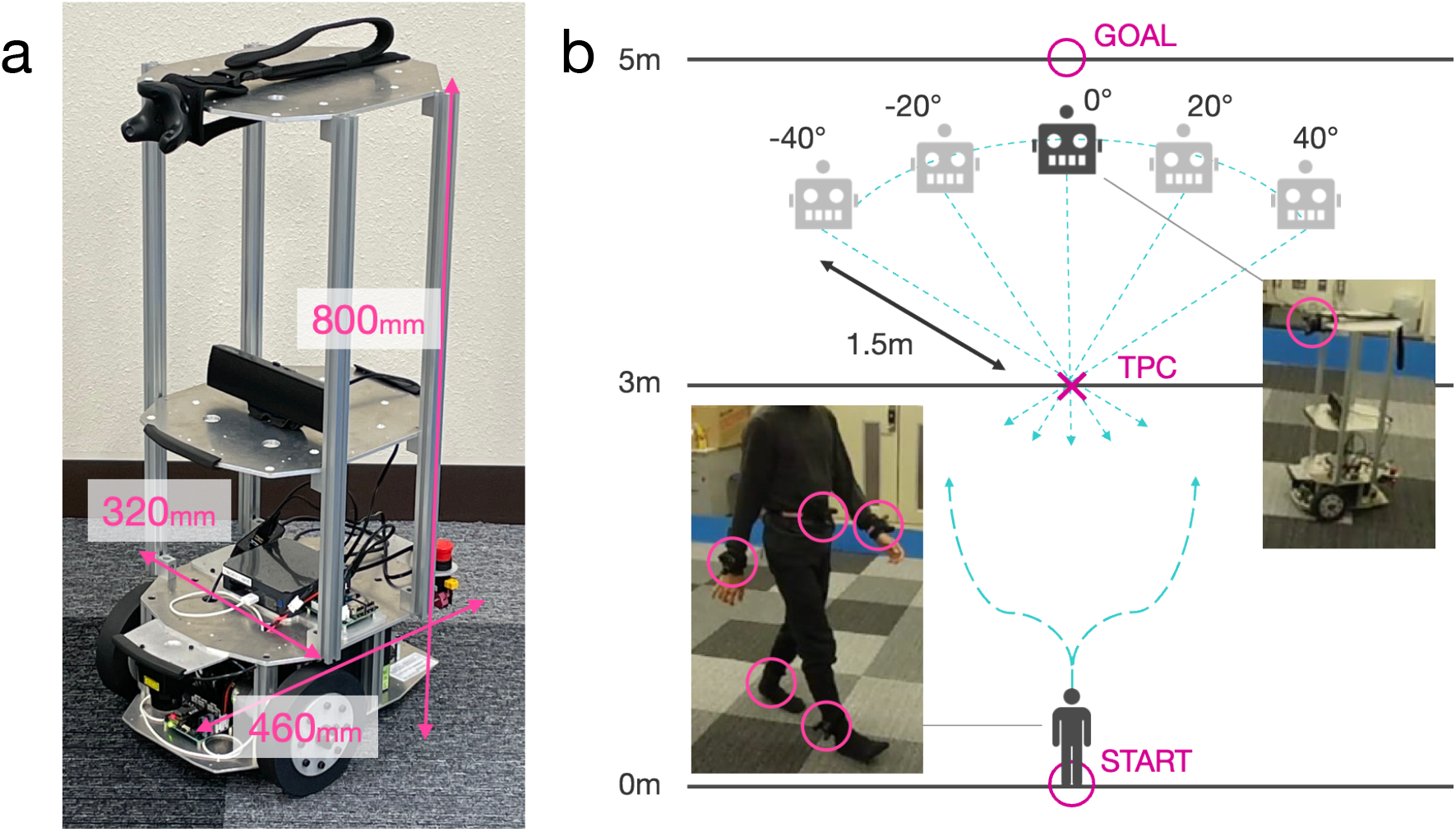
Experimental setup. (a) Autonomous mobile robot (AMR) in the experiment. (b) Illustration of the environment. The participant moves from the starting point toward the goal. The ratio of distances used in this panel differs from the actual ratio used in the study.

##### Tracking system

To measure their motions, the participants were directed to wear five motion trackers (VIVE Tracker 3.0, HTC) on their waists, wrists, and ankles. The AMR had one tracker at the top of its front (**Figures 1a and 1b**). A computer tracking system (with Windows 10) communicated with the trackers controlled by Unity (2020.3.20f1), Steam VR (1.20.4), and Steam VR Unity Plugin (v2.7.3; SDK 1.14.15). The system detected the positions and rotations of each body part with a 90 Hz sampling frequency in a play area defined by four base stations (Steam VR Base Station 2.0) in the experiment room.

##### Environment

**Figure 1b** illustrates the experimental environment of this study. The goal position was located 5 m on a straight line from the starting position of the participants. The theoretical points of collision (TPC) with the AMR were set at the center of the experimental environment and 3 m from the starting point of the participants. The initial positions of the AMR were 1.5 m from the TPC, with five different angles (–40°, –20°, 0°, 20°, and 40°). The field of the experimental environment contained only one participant at a time. Additionally, we only used one AMR for the experiment. The experimenter operated the control and tracking systems from outside this field.

### Procedure and task

First, the AMR randomly traveled to one of the five initial positions. Then, the speaker in the AMR buzzed as a cue to commence the trial. Subsequently, the participant began to walk from the starting point to the goal at a comfortable speed. As the participant took a step, the AMR moved straight toward the TPC at a constant speed of 0.5 m/s. We decided on this speed based on the outcomes of a preliminary experiment to adhere to one of the experimental protocols, “do not let the participants feel scared of the robot.” The end of one trial was declared once the AMR covered a distance of 3 m from its starting position; thereafter, the participant and AMR returned to their initial positions.

The participants were asked to move right or left while walking to evade the AMR. Regarding the technique to avoid collisions, they were instructed to 1) not use extreme evasive actions that could result in an inefficient route toward the goal, 2) not feint an acute change in their course, and 3) evade with a comfortable speed as they would in real life. These instructions were centered on the participants’ natural gait in a corridor or a walkway, where they would not be expected to collide into an AMR. Before the experiment, we informed them about 1) the possibility of an AMR approaching from any of its initial points, 2) the sequence of one trial, and 3) the number of trials in the experiment. No other instructions were provided that could affect their behavior. We executed 50 trials (ten trials × five angles) in a random order for each participant.

### Data analysis

For the analysis, we defined an interval’s beginning as the time at which the AMR sounded the alarm and its ending as the time at which the participant completed walking 5 m in a straight line (or closest to the goal point). We excluded the trials exceeding 10 s as artifacts (corresponding to 1.5 % of all trials) and two trials in which the trackers provided anomalous readings due to a hardware issue. The average trial time for the participants was 7.07 ± 0.20 s. Subsequently, we normalized the trial time from seconds to a percentage value (0%–100%).

In each trial, we estimated the walking trajectory using the three-dimensional (3D) coordinates of the participants’ heads in the space covered by the tracker on their waists. Next, we computed the probability of rightward avoidance. The criterion was the direction that corresponded with a larger change in the mediolateral (ML) displacement at the TPC (i.e., the position reached by the participants after walking straight for 3 m). The average walking speed was estimated from the distance and time of the interval, in which the participants walked between 1 and 3 m in the anteroposterior (AP) direction, to exclude acceleration periods (Souza Silva et al., 2018).

The Palamedes toolbox was used to fit a psychometric function for the probabilities of rightward avoidance at five angles, with the threshold and slope as the free parameters (Prins, 2019; Prins & Kingdom, 2018). When these probabilities formed a step function, we fit them with only the threshold as the free parameter (corresponding to four participants). We excluded one participant from the analysis because we could not fit the psychometric function appropriately as it exhibited an opposite trend. Thus, the data of the remaining fifteen participants were used for further analyses. To quantitatively estimate the participants’ bias for an ML direction, we computed the point of subjective equality (PSE) in each case, as an index of their bias for circumventing to the right or left side.

For motion analyses, we compared the 1) rotation angle and 2) ML displacement using the waist tracker. First, we computed the changes in these two indices for each avoidance direction over a time course. Next, we quantitatively compared the onset times of deviation from the straight line to either side. Specifically, the onset times were defined as the instant at which the waist is twisted over 13.6° (Bruening et al., 2015) and undergoes an ML displacement of more than 0.25 m (Aravind & Lamontagne, 2014; Souza Silva et al., 2018). Please refer to *Motion Analysis* in the **Results** section.

## Results

### Human collision-avoidance behavior

**Figure 2** illustrates two profiles: 1) the average walking velocity and 2) the average distance between the participant and AMR, with each participant’s data. This indicates that the participants accelerate their pace in the first 20 % of the time and then walk steadily at approximately 1.0 m/s between 30 % and 80 % (**Figure 2a**). In addition, the distance was minimized by 62 % on average (**Figure 2b**).

**Figure 2.**
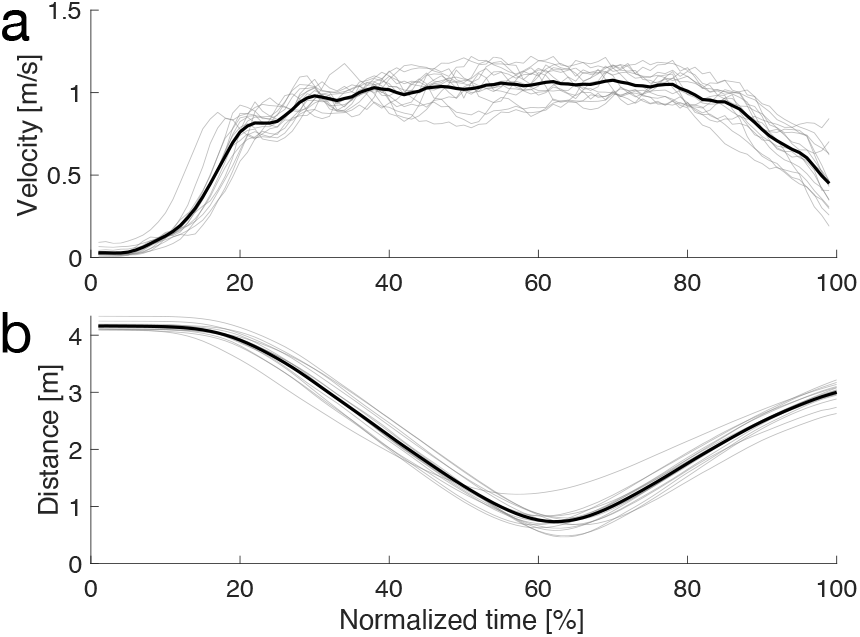
Walking velocity and distance to the AMR. (a) Average velocity profile (thick) and that for each participant (thin). The horizontal axis represents the normalized time between the beginning and end of the trials. The vertical axis represents the velocity. (b) Average distance to the AMR. The other formats are the same as those for (a).

**Figure 3a** illustrates the average walking trajectory in each avoidance direction. Participants in all the trials successfully evaded the AMR, circumventing symmetrically and maintaining a distance from the TPC. Specifically, the participants began circumventing approximately 1.5 m in the AP direction (after a lapse of 4.38 ± 0.27 s). The participants’ average walking speed was 1.03 ± 0.07 m/s, and the minimum distance to the AMR was 0.66 ± 0.18 m.

**Figure 3.**
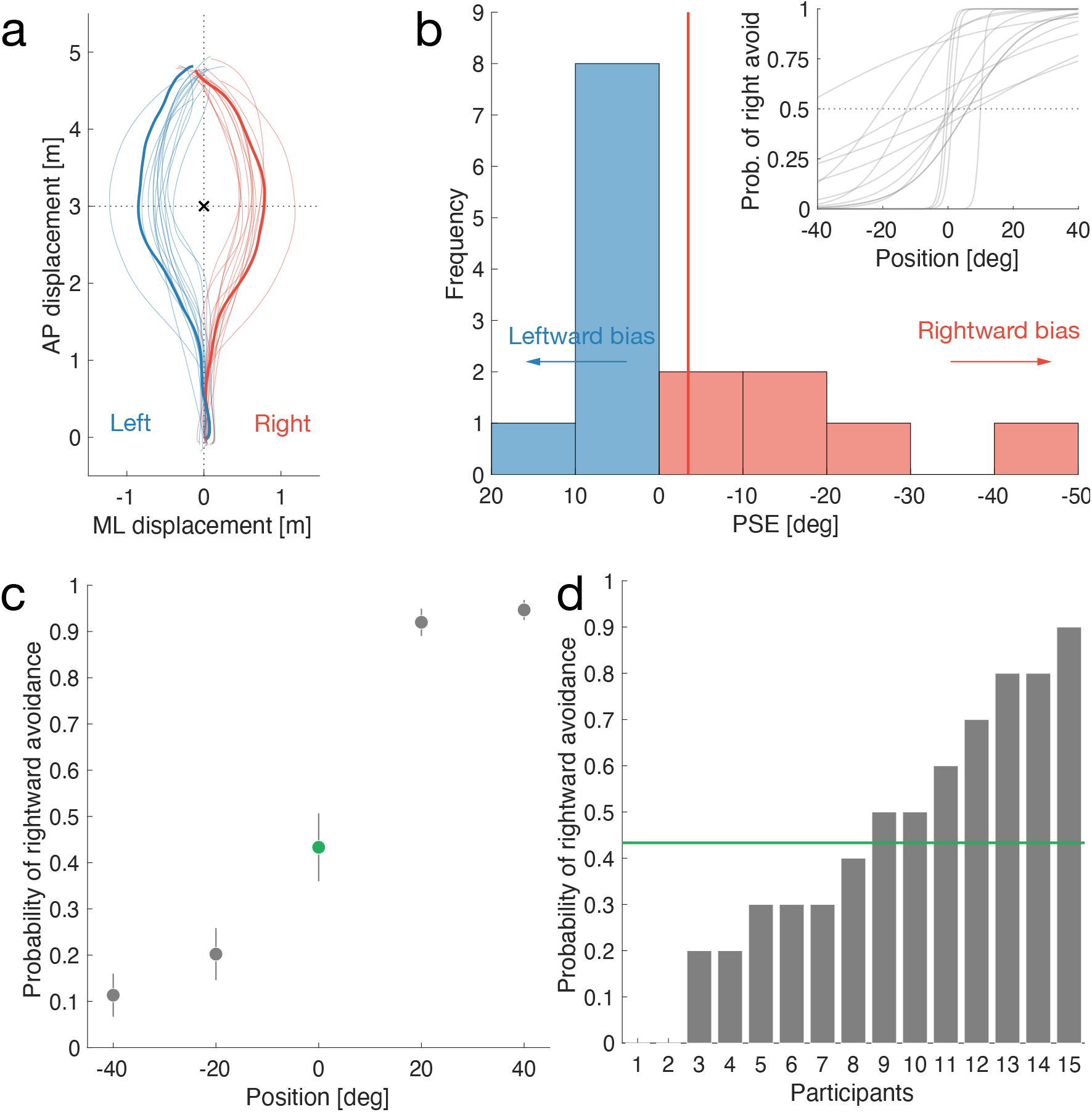
Human collision-avoidance behavior. (a) Walking trajectory. The horizontal and vertical axes represent the mediolateral (ML) and anteroposterior (AP) displacements, respectively. The lines represent the walking trajectories (red: rightward avoidance, blue: leftward avoidance) on average (thick) and for each participant (thin). The black cross represents the location of the theoretical point of collision (TPC). (b) Histogram of the points of subjective equality (PSE) among the participants. The horizontal axis represents the PSE; negative values indicate rightward avoidance. The numbers of participants with rightward and leftward avoidance are six (red) and nine (blue), respectively. The red vertical line represents the average PSE (−3.47°). The image in the top-right corner illustrates the fitted individual curves to estimate the PSEs. (c) Average probability of rightward avoidance (the vertical axis) at the AMR’s initial positions (the horizontal axis). Error bars represent the standard error of the mean. (d) Probabilities of rightward avoidance for each participant (the horizontal axis) in the head-on direction. The vertical axis is the same as that in (c). The green horizontal line represents the average (the same as 0° in (c)).

**Figure 3b** depicts the distribution of the PSE, indicating the directions in which the participants move with respect to the fitted individual curves. The average PSE was –3.47±14.43° (vertical red line in **Figure 3b**), and no significant difference from the origin was observed (*t*(14) = – 0.93, *p* = 0.37, CI = [–11.46, 4.52]). Specifically, the nine left-evading participants had smaller absolute values of PSE and variances compared with the six right-evading persons. Thus, these findings suggest that the participants did not have any preferences regarding the ML direction. In other words, different participants chose different avoidance directions with no common pattern.

A similar “no-bias” trend was observed when the AMR approached the participants head-on (0°; green circle in **Figure 3c**), showing that the average probability of rightward evasion was 0.43 ± 0.28. However, this was composed of avoidance directions with a considerable variance depending on the participants (**Figure 3d**). This implies that, when humans and AMRs cross paths, the precise avoidance direction is difficult to predict using position data alone.

### Motion analysis

We attempted to identify the indicators for predicting the avoidance direction based on human behavior. If the direction can be predicted in advance and if this knowledge was encoded in AMRs, humans could evade robots in a safer, smoother, and more comfortable manner. Hence, we propose a method to predict the avoidance direction based on the positions and rotations measured by the trackers. We assumed that humans twist their waists in the preferred direction immediately before they decide to evade. Therefore, we focused on the waist angle as an apparent cue, which can be measured using common sensors.

**Figure 4a** illustrates that the participants’ average rotation angle (top) peaked faster than the ML displacement (bottom). This finding indicates that the waist sufficiently twists toward its target direction before the rest of the body moves to evade an inbound vehicle. **Figures 4b** and **4c** demonstrate that the participants’ waists twisted significantly before their entire body moved (left: *t*(14) = 4.00, *p* < 0.005, CI = [4.29, 14.21], *d* = 1.03; right: *t*(14) = 3.53, *p* < 0.005, CI = [1.67, 6.83], *d* = 0.91). Thus, before moving, humans involuntarily twist their waists toward the avoidance direction.

**Figure 4.**
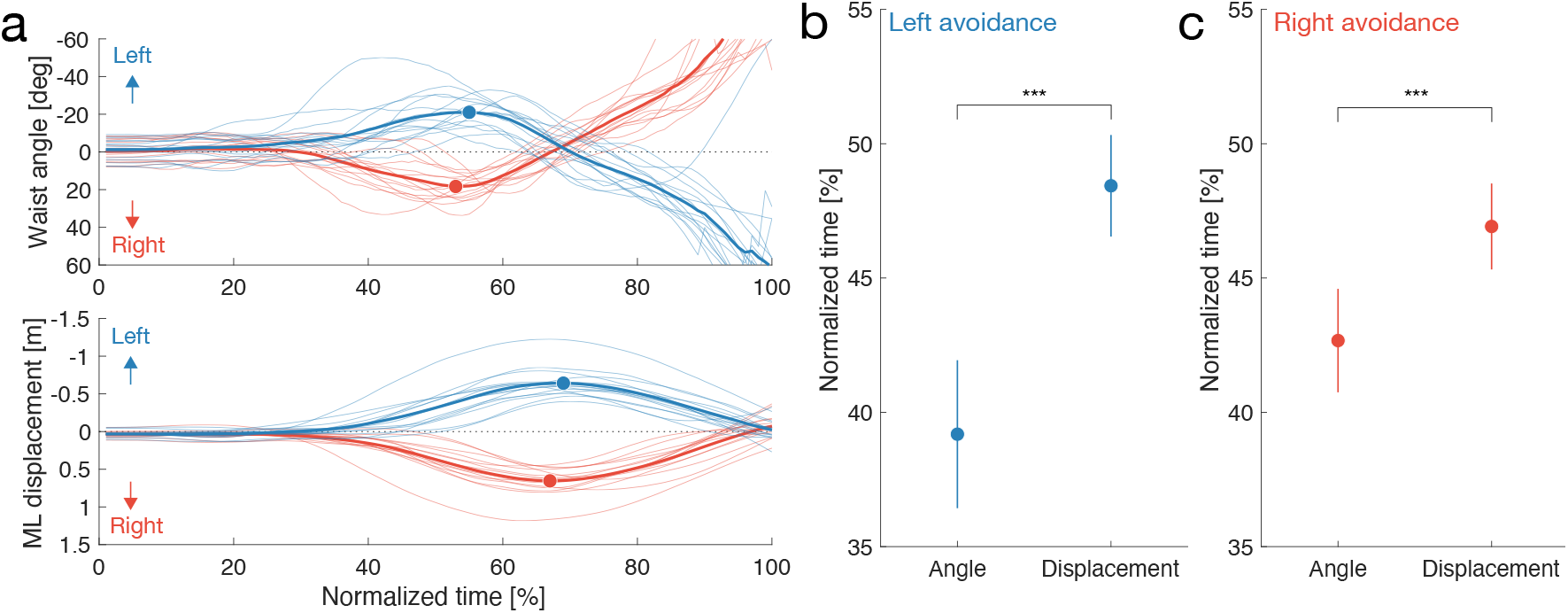
Motion analysis. (a) Changes in the rotation angle and ML displacement at the waist. The horizontal axis indicates the normalized time between the beginning and end of the trials. The vertical axes indicate the waist rotation angle (the top) and ML displacement (the bottom). The other format is the same as that in **Figure 3a**. (b) Comparison of the onset times between rotation angle and ML displacement in the case of leftward avoidance. Error bars represent the standard error of the mean. (c) This panel is the same as (b), except for the avoidance direction. Asterisks indicate statistically significant differences in the *t*-test; *** *p* < 0.005.

## Discussion

This study tested the technique employed by humans to evade an AMR approaching from several directions by recording their motions. As shown in **Figure 3a**, the participants’ walking trajectories represented the same trends as those reported in extant literature (Bühler & Lamontagne, 2019; Souza Silva et al., 2018). This finding suggests that humans’ avoidance trajectory is robust, regardless of the target attribute. By contrast, the average walking speed was marginally slower (1.03 m/s) than that reported in literature. This suggests that humans exercise caution to reduce the risk of collisions (Herman et al., 2005). We did not collect the reference data of the participants’ comfortable speed without the AMR. Meanwhile, the minimum distance to the AMR was shorter (0.66 m). One interpretation is that the participants were solely influenced by the interferer’s (AMR) size because it was smaller than them (Bourgaize et al., 2020). An alternative perspective is that such movements of the AMR could compel humans to approach closer to it due to the real environment in which we accurately estimate a safe distance to avoid a physical collision. For example, a previous study reported that the minimum distance between the participant and the obstacle was shorter in the physical (real) world than in the virtual one (Bühler & Lamontagne, 2019).

We found no significant bias on either side, and the PSEs varied for each participant based on the ease of evasion (**Figure 3b**). Souza Silva et al. suggested that the participants’ bias for rightward avoidance was influenced by the right-handed driving tradition of the area of residence (Souza Silva et al., 2018). By contrast, our results did not indicate such a bias, although participants were marginally more inclined to move leftward (the probability of rightward direction was 43 %) affected by the driving tradition. Hence, this finding cannot be generalized because various factors, including social rules, may determine the evasion strategy.

In addition, our findings were similar to those of a previous study on humans’ evasion of a cylindrical object in a virtual environment (Souza Silva et al., 2018). In other words, the comparison of different targets (humans and the AMR) indicated that the human collision-avoidance strategy differed among humans and with the target object (Vassallo et al., 2017). This suggests that humans employ the object-avoidance strategy against AMRs. More important, our findings indicate that predicting the avoidance direction of humans is difficult when an AMR is approaching head-on.

The participants twisted their waists before the ML displacement (**Figures 4a** and **4b**), suggesting that humans involuntarily twist their waists in the direction of evasion. This indicator of the human body could serve as a factor in determining the avoidance direction. For example, in previous studies, a change in the human body’s center of mass indicated that humans walk using different coordination strategies between steering and circumventing when crossing an obstacle (Vallis & McFadyen, 2003). Studies have also found that involuntary waist rotations by the hanger reflex altered the walking direction (Kon et al., 2016, 2017). This implies that twisting the waist before deciding to move is crucial for determining the walking direction without considering intention or spontaneous action. Thus, we emphasize the importance of focusing on the angle of the waist.

Similarly, studies have revealed that a signal from a body part, such as a gaze, serves as an indicator to predict the avoidance direction (Joshi et al., 2021; Nummenmaa et al., 2009; Saeedpour-Parizi et al., 2021). For example, the gazing direction of a person who approaches head-on affects his/her trajectory (Nummenmaa et al., 2009). This implies that, although humans guess the direction of a person based on the tilt of his/her head, they gaze at the facing person as a signal to indicate their directions, and vice versa. Similarly, humans’ shoulder rotation becomes an indicator of the passability of apertures (Hackney et al., 2015). However, head rotation is not associated with the walking direction (Cinelli & Warren, 2012). Furthermore, a previous study reported no significant difference between the mean onset of the ML deviation and the trunk angle of human walkers as they crossed obstacles (Vallis & McFadyen, 2003). We speculate that body parts nearer to the legs, such as the waist, are more closely associated with the waking direction. Overall, our findings suggest that the avoidance direction of humans can be predicted based on their body’s direction, even before they move.

Although we only focused on the waist movement to predict the avoidance direction, the avoidance direction can be more precisely predicted by analyzing other motions such as the movements of the wrists and ankles. For example, motion and gait features serve as indicators for recognizing Parkinson’s disease (Ťupa et al., 2015). A recent study reported that a person’s landing foot during walking indicated the predictability of their avoidance direction (Tomizawa & Shibata, 2016). Thus, a difference in the avoidance direction derived from the individuals could be determined by various factors (i.e., each part of the body).

Further studies using additional signals would solve these challenges. The technique by and context in which humans employ evasive actions and navigate must be explored. Based on the findings, we conclude that humans and AMRs could cross paths more smoothly if the latter can detect the rotation angle of the waist. Furthermore, the avoidance probability or distance will change when more “human-like” features are provided to the AMR. For example, although this study used a simple wheeled AMR, changing its appearance to emulate that of a human would positively influence pedestrian behavior. Similarly, in terms of actions, the AMR could “bow” when crossing paths with an individual to elicit a human-to-human-like behavior from the individual. Thus, these approaches could render human–AMR interactions friendlier and more organic.

Notably, this study involves a few limitations. First, we did not consider the experimental conditions for the crossing sequence of the participants and AMR or provide relevant instructions. Silva et al. proposed an effective measure based on robotics that focused on boundary dividing collision avoidance into “first” or “subsequent” between humans and robots (Silva et al., 2019). Comparing this to approach more directly with the paradigm of this experiment, the participants would be explicitly instructed to evade either before or after the TPC. Second, although the AMR’s speed was set to 0.5 m/s, humans change their responses with variations in the AMR’s speed. Third, to better mimic real-world scenarios, the participants should interact with the AMR after they have reached a steady-state self-selected walking speed. This suggests that a larger space or an environment wherein multiple objects can sufficiently interact would be more effective, thus bridging the gap between the laboratory and real-world conditions.

## Conclusion

In summary, we clarified the avoidance direction of humans against a passing AMR. We found no significant bias among humans toward rightward or leftward avoidance. We suggest employing a strategy that views the AMR as an approaching object rather than a human. In addition, the findings revealed that humans twist their waists before turning to the direction in which they wish to move. Further studies are required to identify other indicators of the human body against approaching AMRs to ensure safer environments. Such findings would support the development of AMRs for smart cities.

## Acknowledgements

This paper is based on results obtained from the project JPNP20004, which was subsidized by the New Energy and Industrial Technology Development Organization (NEDO). This work was supported by the CASIO SCIENCE PROMOTION FOUNDATION (39-55) and the 2021 Toyohashi University of Technology President Funding (Young Researchers).

## Data availability

The datasets generated during and/or analyzed during the current study are available from the corresponding author on reasonable request.

